# Haplotype-phased, chromosome-level, annotated genome assembly of the soybean rust pathogen *Phakopsora pachyrhizi*

**DOI:** 10.64898/2026.09.29.755439

**Authors:** Nicholas Greatens, Manjula G. Elmore, Sowmya R. Ramachandran, Amy Wright, Luke J. Tallon, Steven A. Whitham, Rachel A. Koch Bach, Kerry F. Pedley

## Abstract

*Phakopsora pachyrhizi*, the causal agent of soybean rust, is among the most aggressive and economically impactful pathogens affecting soybean (*Glycine max*) production worldwide. As a dikaryotic rust fungus, *P. pachyrhizi* maintains its genetic information as two discrete haplotypes that are sequestered within separate nuclei for most of its life cycle as it reproduces asexually via the production of infectious urediniospores. Previous genomes of three *P. pachyrhizi* isolates collected in South America revealed a large genome with a high transposable element content, measuring up to 1.25 Gb and 93%, respectively. Although these efforts provided valuable foundational resources, the complexity of the genome resulted in highly fragmented assemblies. Here, we generated a highly contiguous, substantially improved, annotated genome resource from the *P. pachyrhizi* isolate LA04-1, collected during the initial outbreak of soybean rust in the continental United States in 2004. The assembly is haplotype-phased, with each haplotype assembled into 18 chromosomes totaling 572 Mb and 569 Mb in size. The isolate contains unique mating-type genes and does not share haplotype genomes with any available isolates with whole-genome sequencing datasets. This resource will be valuable to future investigations into the underlying mechanisms of virulence in this important plant pathogen.

## Introduction

Soybean (*Glycine max*), grown for its high protein and oil content, is one of the most important commodity crops in the world. Among the most economically damaging diseases affecting soybean production is soybean rust, caused by the fungal pathogen *Phakopsora pachyrhizi* (Yorinori et al. 2005; Murithi et al. 2016). Originally described in Japan over a century ago (Hennings 1903), *P. pachyrhizi* is now present in all major soybean growing regions (Kelly et al. 2015), including the United States (Schneider et al. 2005).

Under high disease pressure and environmental conditions that favor disease, such as in tropical and subtropical climates, *P. pachyrhizi* can cause yield losses up to 90% (Kelly et al. 2015).

*Phakopsora pachyrhizi* belongs to the order Pucciniales, or the rust fungi, a large taxon of obligately biotrophic, economically important plant pathogens. Rust fungi have complex and variable life cycles with up to five distinct spore types and may be heteroecious, requiring a secondary host to complete sexual reproduction (Kolmer et al. 2009). On soybean and other susceptible legume hosts, *P. pachyrhizi* reproduces asexually through rust-colored urediniospores. Although another spore type, the teliospore, has been observed on several legume hosts, teliospore germination and basidiospore formation have been documented only under laboratory conditions (Goellner et al. 2010). The complete life cycle is unknown, and neither sexual reproductive structures nor alternate hosts have been reported (Goellner et al. 2010). Nevertheless, two lineages of *P. pachyrhizi* have been described based on whole genome SNP data (da Rocha et al. 2025): one genetically diverse lineage composed of four isolates collected between 1972–1994 in Australia, Hawaii, India, and Taiwan; and a second, widespread, clonal lineage comprised of recently collected isolates in South America, Africa, and Asia. Recent population genetics studies have shown that two populations are now present in the continental United States (Clary et al. 2026), and that isolates of soybean rust collected in the continental United States from 2007–2008 are genetically distinct from South American and East African populations (Ferreira et al. 2025).

The first published genome assemblies of *P. pachyrhizi* (Gupta et al. 2023) were generated from three purified isolates collected in South America and revealed a large genome (1.1 to 1.3 Gb, including both haplotypes) rich in transposable elements (TEs). Assemblies provided valuable insights into TE evolution, variable expression patterns among the three clonal isolates, and pathogen effectors—small, secreted proteins that may confer either virulence or resistance, depending on the host genotype (Cui et al. 2015). While the presence of expected BUSCO genes indicates the assemblies and annotations are high quality, assemblies are unphased and fragmented. Here, we announce the genome of *P. pachyrhizi* isolate LA04-1. The assembly is highly contiguous, assembled in 36 chromosomal scaffolds phased into two haplotype genomes of 572 Mb and 569 Mb, each with 18 chromosomes. Haplotype assemblies and homeodomain (HD) transcription factor encoding mating type (MAT-1) alleles of isolate LA04-1 are unique to the lineage, based on k-mer containment. This genomic resource will be valuable to illuminate the evolution and diversity of *P. pachyrhizi* and in investigations into host resistance and virulence.

## Materials and methods

### Sample collection and sequencing

The *P. pachyrhizi* isolate LA04-1 was collected in Ben Hur, Louisiana in 2004 and increased via four successive rounds of single pustule purification on the susceptible soybean cultivar ‘Williams’ as previously described (Pham et al. 2009). It has since been further propagated on the susceptible soybean cultivar ‘Williams 82’. Urediniospores were collected using a mechanical harvester and stored under liquid nitrogen at the USDA-ARS Foreign Disease-Weed Science Research Unit biological safety level 3 (BSL-3) pathogen containment facility at Fort Detrick, Maryland, USA.

High-molecular weight DNA from 500 mg of LA04-1 urediniospores was extracted following the protocol developed by (Schwessinger and Rathjen 2017), purified with an NEB Monarch Spin gDNA Extraction Kit (New England Biolabs, Ipswich, MA), and sheared to an average size of 15kb on a Hamilton Microlab Starlet instrument (Hamilton, Reno, NV). The library was constructed with a SMRTbell Prep Kit 3.0 (Pacific Biosciences, Menlo Park, CA) per manufacturer protocol, size-selected for fragments greater than 10kb with a BluePippin instrument (Sage Science, Beverly, MA), and sequenced with SPRQ chemistry on a PacBio Revio instrument (Pacific Biosciences, Menlo Park, CA) following standard protocol by Maryland Genomics at the University of Maryland Institute for Genome Sciences (Baltimore, Maryland, USA). Hi-C data was generated by cross-linking DNA with a Proximo Hi-C 5.0 Kit (Phase Genomics, Seattle, WA) following the manufacturer’s protocol and sequenced with an Illumina NovaSeq X Plus (San Diego, California, USA) by Maryland

Genomics. Short reads were generated by germinating 10mg of urediniospores in 50mL water amended with ampicillin (50 µg/mL). After 16h, germinated spores were collected, blotted with a Kimwipe, and homogenized with a Qiagen TissueLyser II (Qiagen, Valencia, CA) at 30 Hz for 60 seconds in 2 mL Eppendorf Safe-Lock tubes containing 2 mm zirconium oxide beads (Next Advance, Troy, NY, USA). DNA was extracted with an Omega Biotek E.Z.N.A.® HP Plant & Fungal DNA Kit (#D2485; Norcross, GA, USA) and sequenced by AmpSeq (Gaithersburg, Maryland, USA) with an AVITI machine (Element Biosciences, San Diego, CA, USA).

RNA-Seq data was generated for three biological replicates each of resting spores, germinated spores, and appressoria from *P. pachyrhizi* isolate LA04-1 (TABLE S1). Germinated urediniospores were prepared as described above, and appressoria were prepared as described by Stone et al. (2012). In brief, 150 mL of a suspension containing 100,000 urediniospores/ml was applied to the surface of a 245 mm × 245 mm × 20 mm polystyrene dish and incubated in the dark at room temperature for 6 h. A sterile water wash was used to remove any ungerminated urediniospores, and plates were examined under a microscope before and after washing to confirm the presence of appressoria.

Appressoria were collected by scraping the plate surface with a cell scraper (Nunc, Thermo Fisher Scientific; Waltham, MA) and immediately frozen and stored in liquid nitrogen.

Resting urediniospores, germinated urediniospores, and appressoria were each separately ground under liquid nitrogen with a mortar and pestle. Total RNA was extracted using the Qiagen RNeasy Plant Midi Kit (Qiagen, Valencia, CA) and mRNA was isolated using the Dynabeads mRNA Direct Kit (Invitrogen, Thermo Fisher Scientific; Waltham, MA). Libraries were prepared with the NEBNext Ultra II Directional RNA Library Prep Kit (New England Biolabs, Ipswich, MA). Three libraries per group were sequenced with 150 bp paired end sequencing on an Illumina NovaSeq6000. In addition, the three biological replicates from each group were pooled into a single library prepared with the same kit and sequenced with 250 bp paired-end sequencing on an Illumina NovaSeq6000. All sequencing data were uploaded to NCBI under BioProject PRJNA1518829.

### Assembly

PacBio HiFi reads were screened for quality with NanoPlot 1.42.0 (De Coster and Rademakers 2023), for adapter contamination with HiFiAdapterFilt 3.0 (Sim et al. 2022) and down-sampled to 30x coverage per haplotype with FiltLong 0.2.1 (github.com/rrwick/Filtlong), retaining the longest reads with an average quality greater than 20. Hi-C reads were preprocessed with fastp 0.23.4 (Chen et al. 2018) using default settings. Hifiasm 0.25.0 (Cheng et al. 2021) was used to assemble the genome with Hi-C reads for phasing. Contigs supported by at least 15x average coverage were retained.

Coverage-filtered contigs were queried against the published *P. pachyrhizi* mitogenome (NC_014344.1; Stone et al. 2010) with BLASTn 2.15.0 megablast (Camacho et al. 2009), and reads supporting 34 contigs aligning to the mitochondrial reference were reassembled with Flye 2.9.4 (Kolmogorov et al. 2019) into a single contig, and the resulting mitogenome assembly was oriented on a gene with Circlator 1.5.5 (Hunt et al. 2015). Remaining contigs were assessed for contamination through BLASTn search against the nt database, and six contigs originating from soybean and whitefly were removed from the assembly.

Contigs originating from the nuclear genome from both haplotype assemblies were concatenated, and preprocessed Hi-C reads were mapped to the assembly following recommendations for use with HapHic 1.0.6 (Zeng et al. 2024). Contigs from each haplotype were separately scaffolded with YAHS 1.2.2 (Zhou et al. 2023), and scaffolds were submitted to the HapHic pipeline to generate files for manual curation in Juicebox 2.17.00 (Durand et al. 2016). No phase switches or mis-joins were detected based on visual inspection. The presence of terminal telomeric sequences was assessed with tidk (Brown et al. 2025) and FindTelomeres (github.com/FindTelomeres), and nine chromosomes contained poorly supported, likely erroneous sequences that extended less than 3000 bp past telomeric repeats and were trimmed to the telomere based on minimap2 2.24 (Li 2018) alignments of the filtered HiFi reads to the assembly. Unplaced contigs were removed from the assembly if over 99 % of the sequence aligned to another contig or scaffold and was over 99 % identical in the aligned block. The curated assembly was polished using NextPolish2 0.2.2 (Hu et al. 2024), based on Winnowmap 2.03 (Jain et al. 2020) alignments of the filtered HiFi reads to the assembly and mitogenome and on 21-and 31-mers present in short reads generated from the sample at 60x coverage and prepared with yak v1-r69 (github.com/lh3/yak).

### Annotation

Transposable elements were annotated with the EDTA pipeline 2.2.2 (Ou et al. 2019) and masked with RepeatMasker 4.1.5 (Smit et al. 2013). RNA-Seq reads were aligned with HISAT2 2.2.1 (Kim et al. 2019) to both sets of chromosomes, unplaced contigs, mitogenome, and the soybean ‘Williams 82’ reference assembly, including organellar genomes (GCF_000004515.6; Schmutz et al. 2010). Read pairs with at least one read aligning to *P. pachyrhizi* chromosomes and contigs were realigned with HISAT2 to each haplotype assembly separately. Genes were predicted separately for each haplotype assembly with the BRAKER 3.0.8 (Gabriel et al. 2024) pipeline using the RNA-Seq data aligned separately to each haplotype and the OrthoDB 12.2 Basidiomycota protein dataset (Tegenfeldt et al. 2025). In addition, Funannotate 1.8.16 (Palmer and Stajich 2023) predict was run, utilizing the Augustus model trained on haplotype one from the BRAKER3 annotation, the RNA-Seq alignments, and gene models from PASA 2.5.2 (Haas et al. 2008) based on genome-guided transcript assemblies from StringTie 3.0.3 (Pertea et al. 2015) and *de novo* assemblies with Trinity 2.15.1 (Grabherr et al. 2011) of reads where at least one paired read aligned to *P. pachyrhizi* chromosomes or contigs. Funannotate predictions were updated in two rounds in PASA. In each case, unplaced contigs were annotated with haplotype two.

Gene predictions were evaluated for completeness with BUSCO v6.1.0 (Simão et al. 2015) with the Basidiomycota_odb12.2 dataset (Tegenfeldt et al. 2025). BUSCO scores indicated that BRAKER generated substantially more complete gene predictions and was used as the main source of gene prediction (TABLE S2). Unique genes predicted by the Funannotate pipeline were merged with the BRAKER annotation if they encoded secreted proteins, based on SignalP 6 predictions (Teufel et al. 2022) to improve annotation of candidate secreted effectors. Annotations were merged and processed with the AGAT 1.0.0 (Dainat et al. 2022) scripts “agat_sp_merge_annotations.pl”, “agat_sp_manage_IDs.pl”, “agat_sp_fix_overlaping_genes.pl”, and “agat_sp_fix_features_locations_duplicated.pl”.

Functional annotations were generated with Funannotate annotate with additional input from InterProScan v6.0.2.2 (Blum et al. 2026), Phobius (Käll et al. 2004), and AntiSMASH v8.0.4 (Blin et al. 2023).SignalP 6 and EffectorP 3.0 (Sperschneider and Dodds 2022) were used to predict candidate secreted effector proteins from candidate secreted proteins. The mitochondrial genome was annotated separately with MFannot v1.37 (Lang et al. 2023). Output from both pipelines and the final annotation are available at Ag Data Commons: 10.15482/USDA.ADC/34005444.

### Comparison with references

Three published assemblies of *P. pachyrhizi* were available for comparison at the time of writing (TABLE 1; Gupta et al. 2023), K8108, MT2006, and UFV02. Genome completeness statistics of each assembly were assessed with BUSCO 6.1.0 (Simão et al. 2015) using the Basidiomycota_odb12.2 dataset (Tegenfeldt et al. 2025) and gfastats (Formenti et al. 2022). Predicted proteomes of both haplotypes of LA04-1 and the isolates listed above were submitted to Orthofinder v 3.0.1 (Emms and Kelly 2019). Predicted proteomes were also screened for secretion signals with SignalP as described above, and effectors were predicted with EffectorP. Predicted protein sequences were compared based on predicted sequences using seqkit rmdup (Shen et al. 2024), and based on orthology, from Orthofinder results, available in the Ag Data Commons repository.

**Table 1:**
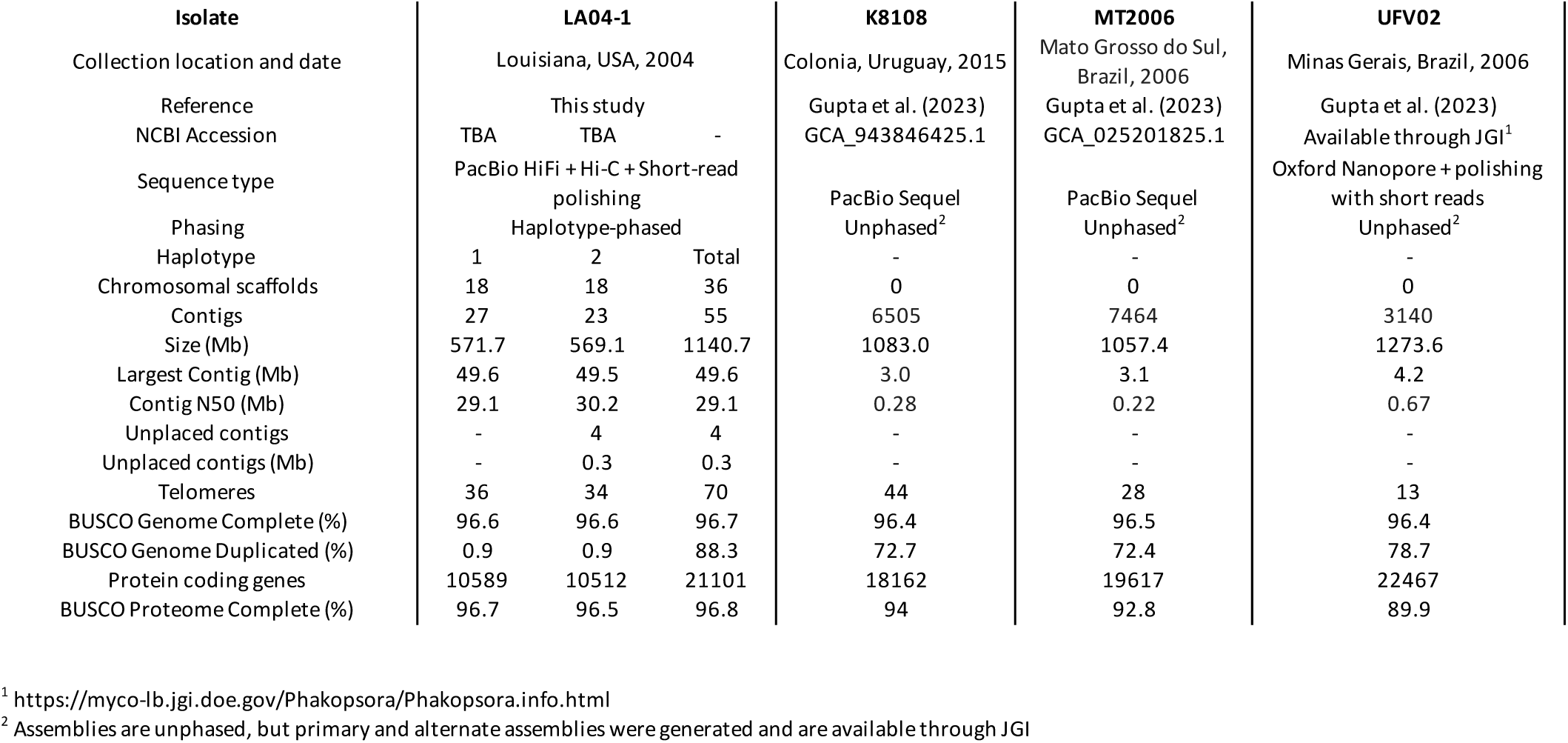
Genome assembly and annotation statistics for P. pachyrhizi LA04-1 and other P. pachyrhizi assemblies

Unassembled short reads from representative isolates of *P. pachyrhizi* clades 1 and 2 (da Rocha et al. 2025), were screened for containment of LA04-1 haplotype assemblies and of mRNA sequences of predicted homeodomain-encoding mating type (MAT-1) alleles with Mash v2.3 (Ondov et al. 2016), following Sperschneider et al. (2023).

## Results and discussion

The genome of *P. pachyrhizi* isolate LA04-1 was assembled in 36 chromosomes and phased with Hi-C contacts (FIG 1A) into two haplotypes of 572 Mb and 569 Mb, each with 18 chromosomes (TABLE 1), a conserved number of chromosomes in most rust fungi (Spanner et al. 2026). Of the 36 total chromosomes, 24 were assembled end-to-end in single contigs, and the remainder were scaffolded from two or three contigs. An additional four contigs, totaling 283 kb in length, were unplaced. Telomeric repeats are present on both ends of 34 chromosomes, while two lack a single telomere, and in each assembly, over 96 % of BUSCO genes are present and annotated. Like the genomes of other rust fungi, the genome is highly repeat rich, with 87 % of the genome composed of transposable elements, or mobile genetic elements (TABLE S3), based on predictions by EDTA. Long Terminal Repeat retrotransposons (LTR-RTs) compose 71.4 % of the genome, and Terminal Inverted Repeat (TIR) transposons compose 10.9 %. Haplotypes are generally syntenic and do not contain any translocations greater than 100 Kb, but several small inversions are present (FIG 1B).

In haplotypes 1 and 2, respectively, and 10589 and 10512 genes were annotated, including 1246 and 1329 genes encoding secreted proteins and 815 and 914 genes encoding candidate secreted effectors (TABLE S2). When considering multiple isoforms, 1734 unique secreted proteins are predicted to be effectors, 454 of which are encoded by both haplotypes. The remainder are predicted to be encoded by one haplotype, due to heterozygosity or hemizygosity, where a gene is present in only one haplotype.

Respectively, haplotypes 1 and 2 encode proteins in 116 and 126 orthogroups without orthologues in the other haplotype.

Isolate LA04-1, collected shortly after the first report of soybean rust disease in the United States, is avirulent to numerous soybean rust resistance genes, making the isolate highly valuable to investigations of host resistance genes, including *Rpp1*, *Rpp2*, *Rpp3* and *Rpp4* (Meyer et al. 2009; Pedley et al. 2019; Bish et al. 2024; Holan et al. 2026). Neither haplotype genome of LA04-1 nor its homeodomain (HD)-encoding MAT-1 mating type genes is shared with other isolates with available short-read whole genome sequencing data, including the isolates within the genetically diverse clade Pp1 or the widespread Pp2 clade, described by da Rocha et al. 2025 (TABLE S4), indicating LA04-1 likely represents a distinct lineage.

Consistent with its genetic distance from other known isolates, a comparison of the predicted proteome of LA04-1 with those of available reference genomes shows the isolate also varies in candidate effector content: of the 1734 total candidate effectors, 1008 proteins and 149 orthogroups are unique to the isolate, based on available gene annotations. Similarly, however, the isolates UFV02, K8108, and MT2006, also vary in their sequences of predicted effector proteins (TABLE S5), despite their clonality. None contains unique predicted effector orthogroups but protein sequences are variable among isolates (TABLE S5). Improved and consistent gene annotation as well as sequence quality across *P. pachyrhizi* isolates and lineages will improve annotation of predicted effectors.

Long-read sequencing technologies combined with Hi-C or similar methods have recently made possible haplotype-phased, chromosome-level assemblies of rust fungi (Henningsen et al. 2025), reshaping our understanding of their evolution. In asexual lineages of cereal rust fungi, for example, genotypic and phenotypic diversity has evolved through various novel mechanisms (Figueroa et al. 2020), including internuclear exchange of chromosomes (Li et al. 2025), somatic exchange of haploid nuclei (Li et al. 2019; Sperschneider et al. 2023; Henningsen et al. 2024; Holden et al. 2025), and chromosomal structural variation including megabase-scale translocations (Spanner et al. 2026). In contrast to the cereal rust fungi, however, few genome resources have been developed for *P. pachyrhizi*, and previously available assemblies represented a single clonal lineage (da Rocha et al. 2025).

In completeness and contiguity, this genome resource improves substantially upon previously available references (TABLE1). As the first phased reference genome of *P. pachyrhizi*, and the first genome for an isolate outside the lineage predominant in South America and Africa, this genomic resource promises to be useful for investigations into the genetic diversity, population structure, gene expression, and virulence of *P. pachyrhizi*, a major pathogen of soybean worldwide. The generation of haplotype-phased genome references for other lineages will be useful to investigate effector diversity among isolates and may enable the association of effector proteins with their corresponding host resistance genes, significantly benefiting research into soybean rust disease resistance and the development of resistant varieties.

## Data availability

All sequencing data is available through NCBI under BioProject PRJNA1518829. Genome assemblies and annotations are available under the NCBI accession numbers TBA and TBA (pending review) and at Ag Data Commons 10.15482/USDA.ADC/34005444. Data can also be accessed at: https://figshare.com/s/7c52266d0eda206db96c.

## Acknowledgments

All opinions expressed in this paper are the authors’ and do not necessarily reflect the policies and views of the USDA, DOE, or ORAU/ORISE. Mention of trade names or commercial products in this publication is solely for the purpose of providing specific information and does not imply recommendation or endorsement by the USDA. The USDA is an equal opportunity provider and employer.

---

**FIG 1.**
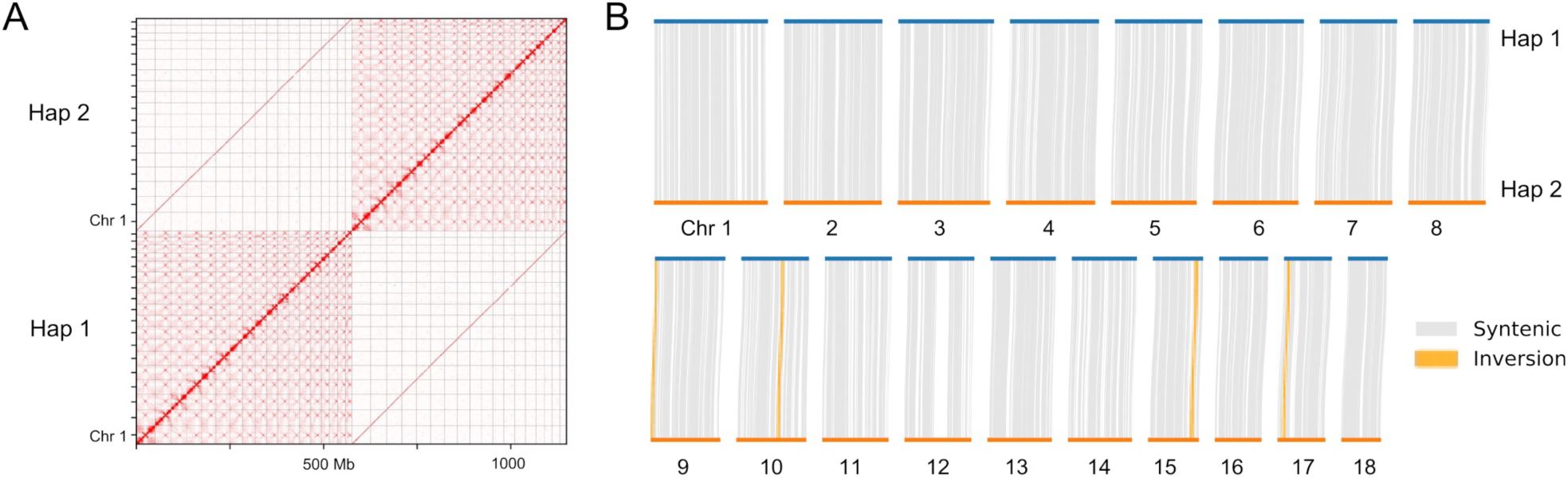
Hi-C contact map and synteny plot for *Phakopsora pachyrhizi* isolate LA04-1 haplotypes one and two. (a) Hi-C contact map; (b) Synteny plot comparing chromosomes. Hi-C contacts phased chromosomal scaffolds into nuclear compartments. Chromosomes are generally syntenic and contain no large translocations. Plots were generated with HapHic 1.0.6 (Zeng et al. 2024) and plotsr 1.2.0 with syri 1.8.3 (Goel et al. 2019; Goel and Schneeberger 2022).

