## Supplementary_tables for "Haplotype-phased, chromosome-level, annotated genome assembly of the soybean rust pathogen *Phakopsora pachyrhizi*"

**Table S1. RNA-Seq data accessions, conditions, and read mapping rates**

| SRA | Condition | Replicate | Reads before filter (Mil.) | Bases before filter (Gb) | Reads after filter (Mil.) | Bases after filter (Gb) | Reads mapped to LA04-1 nuclear genome (Mil.) | Bases mapped to LA04-1 nuclear genome (Gb) | Reads aligned to LA04-1 nuclear genome (%) |
|---|---|---|---|---|---|---|---|---|---|
| SRR40373608 | Resting | 1 | 57.9 | 8.7 | 57.4 | 8.4 | 53.3 | 7.8 | 92.9 |
| SRR40373607 | Resting | 2 | 54.2 | 8.1 | 53.7 | 7.8 | 49.9 | 7.3 | 92.9 |
| SRR40373606 | Resting | 3 | 58.5 | 8.8 | 58.1 | 8.5 | 54 | 7.9 | 92.9 |
| SRR40373605 | Germinated | 1 | 41.3 | 6.2 | 40.8 | 6 | 38.1 | 5.6 | 93.4 |
| SRR40373604 | Germinated | 2 | 50.4 | 7.6 | 49.9 | 7.3 | 46.5 | 6.8 | 93.2 |
| SRR40373603 | Germinated | 3 | 45.9 | 6.9 | 45.4 | 6.7 | 42.5 | 6.3 | 93.6 |
| SRR40373602 | Appressoria | 1 | 46.1 | 6.9 | 45.6 | 6.7 | 42.6 | 6.2 | 93.4 |
| SRR40373614 | Appressoria | 2 | 50.1 | 7.5 | 49.7 | 7.2 | 46 | 6.7 | 92.6 |
| SRR40373613 | Appressoria | 3 | 44.9 | 6.7 | 44.3 | 6.5 | 40.8 | 6 | 92.1 |
| SRR40373612 | Resting | Pooled from 3 biological reps | 166.1 | 41.5 | 163.1 | 34.1 | 149.5 | 31.2 | 91.7 |
| SRR40373611 | Germinated | Pooled from 3 biological reps | 141 | 35.2 | 138.2 | 29.8 | 127.3 | 27.3 | 92.1 |
| SRR40373610 | Appressoria | Pooled from 3 biological reps | 119.7 | 29.9 | 117.2 | 25.6 | 107.6 | 23.4 | 91.8 |

**TABLE S2. Comparison of annotation methods**

| | Haplotype | Genes | Transcripts | BUSCO 6 completeness (Basidiomycota odb12.2) | Predicted secreted protein-encoding genes (SignalP 6) | Predicted secreted proteins including isoforms (SignalP 6) | Predicted secreted effector-encoding genes (SignalP 6 + EffectorP 3) | Predicted secreted effectors, including isoforms (SignalP 6 + EffectorP 3) |
|---|---|---|---|---|---|---|---|---|
| Braker 3 | Hap1 | 10337 | 12270 | 96.7 | 856 | 981 | 492 | 552 |
| | Hap2 | 10147 | 12585 | 96.5 | 842 | 971 | 502 | 544 |
| Funannotate predict + PASA update[1] | Hap1 | 22719 | 31127 | 85.4 | 1109 | 1781 | 734 | 1181 |
| | Hap2 | 26598 | 34816 | 81.1 | 1156 | 1756 | 804 | 1191 |
| Combined[2] | Hap1 | 10592 | 14529 | 96.7 | 1246 | 2526 | 815 | 1600 |
| | Hap2 | 10515 | 14369 | 96.5 | 1329 | 2546 | 914 | 1636 |

[1] Default weights, with Augustus trained with Braker3 on Haplotype 1

[2] BRAKER3 set with unique Funannotate predicted secreted proteins

**TABLE S3. Repetitive elements (EDTA) summary**

| Class | | Count | Masked (Mb) | Masked (%) |
|---|---|---|---|---|
| **Long interspersed nuclear elements (LINEs)** | | **4933** | **4.74** | **0.41** |
| | L1 | 333 | 0.17 | 0.01 |
| | Tad1 | 4600 | 4.57 | 0.40 |
| **Long terminal repeat (LTR) retotransposons** | | **1142019** | **815.25** | **71.46** |
| | Copia | 70097 | 68.59 | 6.01 |
| | Gypsy | 444177 | 359.10 | 31.48 |
| | Unknown | 627745 | 387.56 | 33.97 |
| **Terminal Inverted Repeats (TIRs)** | | **252829** | **123.74** | **10.85** |
| | CACTA | 25009 | 11.09 | 0.97 |
| | IS3EU | 1012 | 1.12 | 0.10 |
| | Mutator | 161774 | 77.04 | 6.75 |
| | PIF_Harbinger | 6888 | 3.38 | 0.30 |
| | Tc1_Mariner | 6641 | 3.55 | 0.31 |
| | hAT | 51505 | 27.57 | 2.42 |
| | Low complexity | 998 | 0.73 | 0.06 |
| **Helitron** | | **3062** | **3.37** | **0.30** |
| **Unclassified repeat fragments** | | **85218** | **38.90** | **3.41** |
| **Total** | | **1489731** | **987.01** | **86.50** |

**TABLE S4. Containment of LA04-1 haplotype genomes and HD-encoding MAT-1 genes in other datasets**

| Isolate name | SRA | Lineage (da Rocha et al. 2025) | JGI accession ID and project number[1] | Mash contaiment of LA04-1 Hap1: Identity (shared hashes) | Mash contaiment of LA04-1 Hap2: Identity (shared hashes) | Contaiment of LA04-1 Hap1 HD CDS: Identity (shared hashes) | | Contaiment of LA04-1 Hap2 HD CDS: Identity (shared hashes) | |
|---|---|---|---|---|---|---|---|---|---|
| | | | | | | *AC7T4X_2807 (HD1)* | *AC7T4X_2806 (HD2)* | *AC7V7A_2733 (HD1)* | *AC7V7A_2732 (HD2)* |
| FL07-1 | SRR10966826 | Pp2 | Pp_8 (1232866) | 0.995744 (91433/100000) | 0.995812 (91563/100000) | 0.844852 (29/1000) | 0.831966 (21/1000) | 0.867054 (50/1000) | 0.719686 (1/1000) |
| TW-72-1 | SRR10967470 | Pp1 | Pp_13 (1232876) | 0.996378 (92663/100000) | 0.996388 (92682/100000) | 0.853596 (36/1000) | 0.831966 (21/1000) | 0.878593 (66/1000) | 0 (0/1000) |
| IN73-1 | SRR10967480 | Pp1 | Pp_14 (1232878) | 0.995879 (91693/100000) | 0.996311 (92532/100000) | 0.891191 (89/1000) | 0.803086 (10/1000) | 0.852452 (35/1000) | 0 (0/1000) |
| AU79-1 | SRR10967488 | Pp1 | Pp_15 (1232880) | 0.997449 (94777/100000) | 0.997454 (94788/100000) | 0.852452 (35/1000) | 0.838902 (25/1000) | 0.901 (112/1000) | 0.789561 (7/1000) |
| HW94-1 | SRR10967505 | Pp1 | Pp_16 (1232882) | 0.997629 (95137/100000) | 0.997517 (94914/100000) | 0.975658 (596/1000) | 0.948439 (329/1000) | 0.919083 (170/1000) | 0.987753 (772/1000) |
| CO04-2 | SRR10967507 | Pp2 | Pp_18 (1232886) | 0.996462 (92828/100000) | 0.996489 (92881/100000) | 0.853596 (36/1000) | 0.915049 (155/1000) | 0.909686 (137/1000) | 0.933928 (238/1000) |

[1] See also Da Rocha et al. 2025 Table S1

**TABLE S5: Comparison of effectors among LA04-1 haplotypes and isolates MT2006, K8108, and UFV02**

| | Unique effectors (by sequence) | Orthogroups | Unique effector orthogroups | Predicted effectors shared with LA04-1 Haplotype 2 (by sequence) | Predicted effectors shared with MT2006 (by sequence) | Predicted effectors shared with K8108 (by sequence) | Predicted effectors shared with UFV02 (by sequence) |
|---|---|---|---|---|---|---|---|
| **LA04-1 Haplotype 1** | 1054 | 789 | 25 | 454 | 140 | 126 | 112 |
| **LA04-1 Haplotype 2** | 1134 | 799 | 36 | - | 139 | 133 | 106 |
| **LA04-1 Both haplotypes** | 1734 | 915 | 149 | - | 168 | 183 | 140 |
| **MT2006** | 651 | 532 | 0 | 140 | - | 142 | 122 |
| **K8108** | 709 | 578 | 0 | 126 | 142 | - | 93 |
| **UFV02** | 740 | 619 | 0 | 112 | 122 | 93 | - |
